# Plasma iron controls neutrophil production and function

**DOI:** 10.1101/2022.03.17.484503

**Authors:** Joe N. Frost, Sarah K. Wideman, Alexandra E. Preston, Megan R Teh, Zhichao Ai, Lihui Wang, Amy Cross, Natasha White, Yavuz Yazicioglu, Michael Bonadonna, Alexander J. Clarke, Andrew E. Armitage, Bruno Galy, Irina A. Udalova, Hal Drakesmith

## Abstract

Low plasma iron (hypoferremia) induced by hepcidin is a conserved inflammatory response that protects against infections but inhibits erythropoiesis. How hypoferremia influences leukocytogenesis is unclear. Using proteomic data, we predicted that neutrophil production would be profoundly more iron-demanding than generation of other white blood cell types. Accordingly in mice, hepcidin-mediated hypoferremia substantially reduced numbers of granulocytes but not monocytes, lymphocytes or dendritic cells. Neutrophil rebound after anti-GR1-induced neutropenia was blunted during hypoferremia, but was rescued by supplemental iron. Similarly, hypoferremia markedly inhibited pharmacologically-stimulated granulopoiesis mediated by GCSF and inflammation-induced accumulation of neutrophils in the spleen and peritoneal cavity. Furthermore, hypoferremia specifically altered neutrophil effector functions, suppressing antibacterial mechanisms but enhancing mitochondrial ROS-dependent NETosis associated with chronic inflammation. Notably, antagonising endogenous hepcidin during acute inflammation enhanced production of neutrophils. We propose plasma iron modulates the profile of innate immunity by controlling monocyte-to-neutrophil ratio and neutrophil activity in a therapeutically targetable system.

## Introduction

Iron is required for cellular biochemistry, supporting processes such as oxidative metabolism, DNA synthesis and epigenetic remodelling (Andreini et al., 2018; Teh et al., 2021). Sequestration of iron from plasma, termed hypoferremia, commonly occurs during the acute phase of infection, driven by inflammatory induction of the iron regulatory hormone, hepcidin (Drakesmith and Prentice, 2012). Hepcidin blocks iron recycling by haemophagocytic macrophages and iron absorption by duodenal enterocytes, suppressing serum iron concentrations (Drakesmith et al., 2015). This response can protect against certain siderophilic bacterial pathogens, constituting an acute “nutritional immune” mechanism, but if prolonged, leads to inflammatory anemia, and can inhibit proliferative lymphocyte responses to immunization and infection (Arezes et al., 2015; Frost et al., 2021; Littwitz-Salomon et al., 2021).

Of particular note, inflammatory hypoferremia during acute infection frequently coincides with a need to remodel haematopoiesis (Boettcher and Manz, 2017), and to support metabolically-demanding innate cell effector functions – both potentially iron-dependent processes. Whilst studies have highlighted that the neutrophil oxidative burst is impaired by iron deficiency, how cellular innate immunity is affected by plasma iron status and extracellular iron availability more broadly remains unclear (Ahluwalia et al., 2004; Cronin et al., 2019; Egeli et al., 1998; Hassan et al., 2016; Kurtoglu et al., 2003; Monteith and Skaar, 2021; Paino et al., 2009; Renassia et al., 2020). Here we show, via pharmacological manipulation of plasma iron availability in mice, that neutrophil production and functionality are highly iron-sensitive processes, thereby providing insights into how physiologically relevant systemic shifts in nutrient availability modulate innate immunity.

## Results

### Neutrophil production is iron-requiring and sensitive to low iron availability

Erythropoiesis consumes ∼25mg iron per day in humans (Muckenthaler et al., 2017). However, the iron needs of leukocytes are unknown. Having previously modelled iron demands during T cell activation (Teh et al., 2021), we applied the same method to a published proteomic resource (Rieckmann et al., 2017) to estimate the iron content of resting human peripheral blood leukocytes. We predict that neutrophils have significantly higher iron content per cell than other leukocytes (1A). Moreover, a larger proportion of the neutrophil and eosinophil proteomes is ‘iron-interacting’ compared to other leukocytes (1A). Using estimates for the number of leukocytes generated per day (Cosgrove et al., 2021) we predict neutrophil production uses ∼100-fold more iron than B-cells, T-cells or monocytes and only 10-fold less iron than is estimated for erythropoiesis (1B) (Muckenthaler et al., 2017). Thus, neutrophil production is likely to be particularly sensitive to changes in serum iron.

To evaluate the sensitivity of leukocyte subsets to physiological variation in plasma iron availability experimentally, we administered a mimetic of the iron regulatory hormone hepcidin – minihepcidin (mHep) – daily for four days to C57BL/6 mice (1C) (Frost et al., 2021). This caused hypoferremia and suppressed endogenous hepatic hepcidin expression (S1A) (Wang and Babitt, 2019). Mice given mHep had fewer neutrophils and eosinophils, but unchanged Ly6C+ and Ly6C-monocyte frequency, in spleen and blood (1D/E, S1D). Splenic basophils, dendritic cells and lymphocyte subsets (S1B, S1C), and bone marrow (BM) B cells (S1C) were also unaffected. In addition, fewer mature neutrophils (CD101+ Ly6G+ (Evrard et al., 2018)) and Siglec F+ eosinophils were present in the bone marrow after mHep-treatment (1F); in contrast, mature BM monocytes (CXCR4-CD115+ (Chong et al., 2016)) were increased in treated mice (1G). BM neutrophil and monocyte production are driven by highly proliferative committed progenitors: Ly6C+ c-kit+ CD115-CXCR4+ granulocytic myeloblasts and Ly6C+ c-kit+ CD115+ CXCR4+ committed monocyte progenitors (cMoPs) respectively (Evrard et al., 2018; Hettinger et al., 2013). Following mHep-treatment, we observed no change in myeloblasts, but increased cMoP numbers (1H) which could be driving the accumulation of mature BM monocytes.

To probe the differential sensitivity of neutrophil and monocyte lineages to low iron, we quantified CD71 expression (transferrin receptor, which mediates cellular iron acquisition) on committed progenitors. Myeloblasts had significantly higher CD71 expression than cMoPs in control mice, consistent with our *in silico* prediction of greater neutrophil iron demand (1I). Moreover, mHep-treatment markedly increased CD71 expression by both progenitor subsets, likely indicating cellular iron deficiency (1L). A modest increase in the proportion of myeloblasts in S phase was observed in mHep-treated animals, potentially indicating an iron-associated cell cycle perturbation that could link to reduced BM neutrophil production (S1E) (Yu et al., 2007). No difference in cMoP cell cycle profile was observed (S1E). Minihepcidin also suppressed erythropoiesis and increased circulating platelet counts (S1F, S1G) as previously described in low iron conditions (Xavier-Ferrucio et al., 2019).

Alongside a lineage-intrinsic dependence of granulopoiesis on iron, low iron could also indirectly influence granulocyte production. Granulocyte colony stimulating factor (G-CSF) and IL-5 are key cell-extrinsic hematopoietic cytokines controlling neutrophil and eosinophil production respectively (Kopf et al., 1996; Lieschke et al., 1994). We found no evidence of reduced serum G-CSF or IL-5 mHep-treated mice (S1H). In fact, G-CSF was elevated in hypoferremic animals, consistent with the reduced peripheral neutrophil frequency (Stark et al., 2005).

Together these data indicate that minihepcidin-induced hypoferremia associates with altered hematopoietic output, particularly reduced circulating frequencies of neutrophils and eosinophils (1J), in line with the predicted lineage-specific iron demands. In contrast, we observed an expansion of monocytes and their precursors in the bone marrow.

### Hepcidin driven hypoferremia impairs enhanced granulopoiesis

We next considered whether acute hypoferremia would affect the response to enhanced granulopoiesis. We administered low dose anti-Gr1 to mice to transiently and selectively deplete mature neutrophil numbers indicated by peripheral blood cell frequencies 24h after antibody treatment (S2A)), followed by 4 daily mHep doses to induce hypoferremia during granulopoietic recovery (2A). Differentiation from committed granulocyte progenitor to mature neutrophil can be defined by decreased C-kit and increased Ly6G expression (S2B) (Riffelmacher et al., 2017). After 4 days of mHep administration, mature Ly6G-high BM marrow neutrophil production was suppressed, while more immature cells accumulated (2B and S2C); blood neutrophil counts were also reduced (S2D).

To test whether mHep-mediated suppression of neutrophil production was directly related to iron limitation, mice were administered ferric ammonium citrate (FAC, an exogenous iron source) concurrently with mHep following anti-GR1 treatment (2C). FAC rescued CD101+ Ly6G+ BM neutrophil numbers in mHep-treated mice and reduced myeloblast CD71 expression, suggesting cellular iron deficiency was ameliorated (2D); FAC also rescued splenic eosinophil numbers (S2E). Therefore, suppression of granulopoiesis by mHep is iron-dependent.

During inflammation, cytokines such as GCSF drive enhanced neutrophil production, termed emergency granulopoiesis (Manz and Boettcher, 2014). GCSF is also used clinically to mobilize neutrophils post-chemotherapy. We injected mice with GCSF/anti-GCSF complex (more potent than GCSF alone (Rubinstein et al., 2013)) to stimulate granulopoiesis, with or without mHep as above (2E). mHep-treated mice were hypoferremic, while mice treated with GCSF-complex alone had elevated serum iron, likely due to granulopoiesis displacing more iron-requiring erythropoiesis (S2F-G). The expansion of blood and splenic Ly6G+ and mature CD101+ Ly6G+ neutrophils in GCSF-complex treated mice was suppressed by mHep (2F-G, S2H). Although total circulating and bone marrow granulocytes were reduced by mHep-treatment (2H, S2I), a greater proportion of blood granulocytes had an immature CXCR4+ phenotype, and the number of CXCR4+ c-kit+ BM progenitors was elevated (S2J-K) mHep-treatment also led to a modest, but significant, accumulation of CXCR4+ c-kit+ BM granulocyte progenitors in S-phase, which may contribute to reduced production (S2L). In contrast, we found that mHep-treatment did not prevent the GCSF-complex-driven increase of monocytes in blood, spleen and bone marrow (2I and S2M). Thus, mHep-mediated granulopoietic suppression is still observed even in the presence of excess exogenous GCSF-complex, further supporting the hypothesis that the suppressive effect is not due to inhibition of inductive signals.

### Functional impairment of neutrophils by plasma iron deficiency

To examine whether *in vivo* iron restriction influenced neutrophil effector functions, we characterised isolated BM neutrophils (S3A) following anti-GR1 depletion/recovery with and without mHep, as above (S2A). Isolated neutrophils from hypoferremic mice displayed blunted PMA-induced ROS production, impaired phagocytosis of fluorescently-labelled *Escherichia coli* and reduced *Staphylococcus aureus* killing (3A). Production of CCL2 and TNF-alpha upon *ex vivo* stimulation with zymosan was also suppressed (3B).

We then evaluated NETosis, an important neutrophil effector function playing a central role in both host defence and immunopathology (Papayannopoulos, 2018). NETosis was absent in unstimulated neutrophils (S3B), but unexpectedly, given the reduced ROS production of PMA-stimulated iron restricted neutrophils (3A), neutrophils from hypoferremic mice showed a heightened propensity to undergo NETosis in response to PMA-ionomycin stimulation (3C). We confirmed the ROS dependence of NETosis in our hands using combined cytoplasmic and mitochondrial ROS inhibitor DPI (Li and Trush, 1998) (3C and S3C). Attempting to reconcile these NETosis measurements with our observations of the reduced ROS production of PMA-stimulated iron restricted neutrophils (3A), we measured NETosis after PMA stimulation alone. Consistent with our PMA-induced ROS measurements and the known catalytic role of iron in NADPH-oxidase (NOX; Sumimoto, 2008) and myeloperoxidase (Davies, 2011) function, iron depleted neutrophils underwent NOX-dependent NETosis less efficiently in response to PMA (3D and S3D).

Whilst PMA drives NOX-dependent ROS production and NETosis, ionomycin induces NETosis dependent on mitochondrial ROS (mitoROS) (Douda et al., 2015). We evaluated whether iron deficiency disrupted mitochondrial function by assessing neutrophil energy metabolism *in vivo* using SCENITH (Single Cell Metabolism by Profiling Translation Inhibition) (Argüello et al., 2020). Mitochondrial-dependent ATP production was reduced in Ly6G+ granulocytes from mHep-treated mice, whilst proportional glucose dependence was unaffected (3E). In parallel to reduced mitochondrial ATP production, iron-restricted neutrophils produced greater amounts of mitoROS (3F) and had an increased propensity to undergo mitoROS-dependent NETosis in response to ionomycin (3G and S3E).

Monopoiesis is relatively resistant to perturbation by hypoferremia (Fig 1G-H). Therefore, we asked whether monocyte function was also preserved. In splenic monocytes from mHep-treated mice, secretion of IL-12, TNF and IL-6 in response to *ex vivo* LPS stimulation was unchanged compared to monocytes from control mice (S3D).

**Figure 1.**
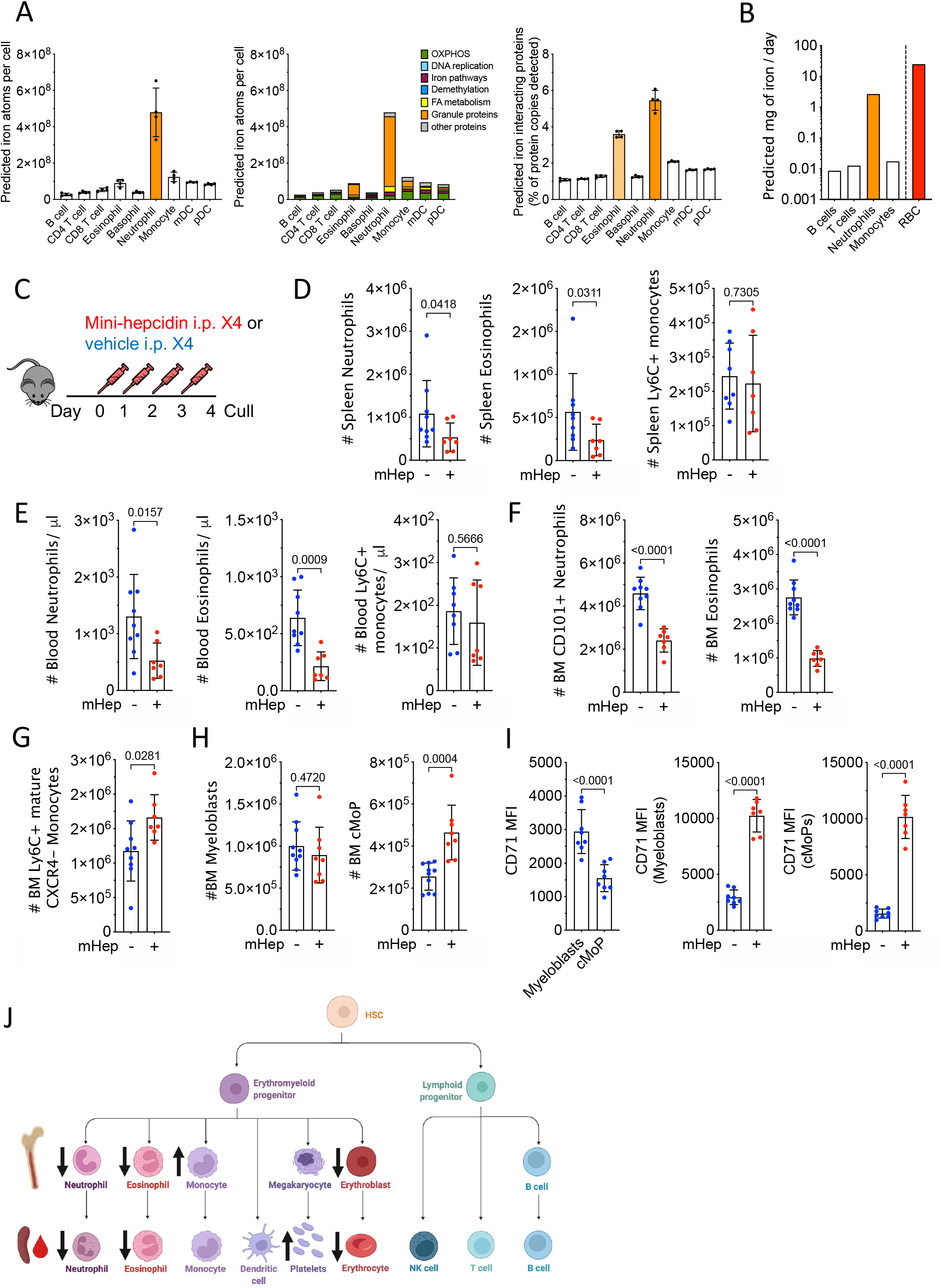
Hepcidin reduces systemic neutrophil numbers. A) Predicted absolute number of iron atoms in total and grouped by pathway, as well as percentage of detected proteins that are iron interacting proteins in resting human peripheral blood leukocytes. Mean ± SD B) Predicted iron requirements (mg / day) for haematopoietic production of the specified lineage, compared to literature-derived value for RBC production. C) Experiment scheme to test effect of mHep on haematopoiesis D) Number of splenic myeloid subsets in experiment described in Fig 1C. Representative of three independent replicates. Neutrophils and Eosinophils Mann-Whitney Test. Monocytes unpaired T-test. Mean ± SD. E) Blood myeloid subsets (numbers / μl) in experiment described in Fig 1C. Representative of three independent replicates. Neutrophils, Welch’s unpaired T-test. Eosinophils and monocytes, unpaired T-test. Mean ± SD. F) Numbers of mature neutrophils and eosinophils in the bone marrow in the experiment described in Fig 1C. Representative of three independent replicates. Unpaired T-test. Mean ± SD. G) Numbers of mature monocytes in the bone marrow in experiment described in Fig 1C. Representative to three independent replicates. Unpaired T-test. Mean ± SD. H) Numbers of committed granulocyte progenitors and committed monocyte progenitors (cMoP) in experiment described in Fig 1C. Representative of three independent replicates. Unpaired T-test. Mean ± SD. I) Comparison of CD71 expression between myeloblasts (granulocyte progenitors) and committed monocyte precursors (cMoP) in experiment described in Fig 1C. Paired T-test. Effect of mHep treatment on CD71 expression by myeloblasts and cMoPs. Unpaired T-test. Mean ± SD. J) Graphical summary of changes in cell numbers observed after minihepcidin treatment.

**Figure 2.**
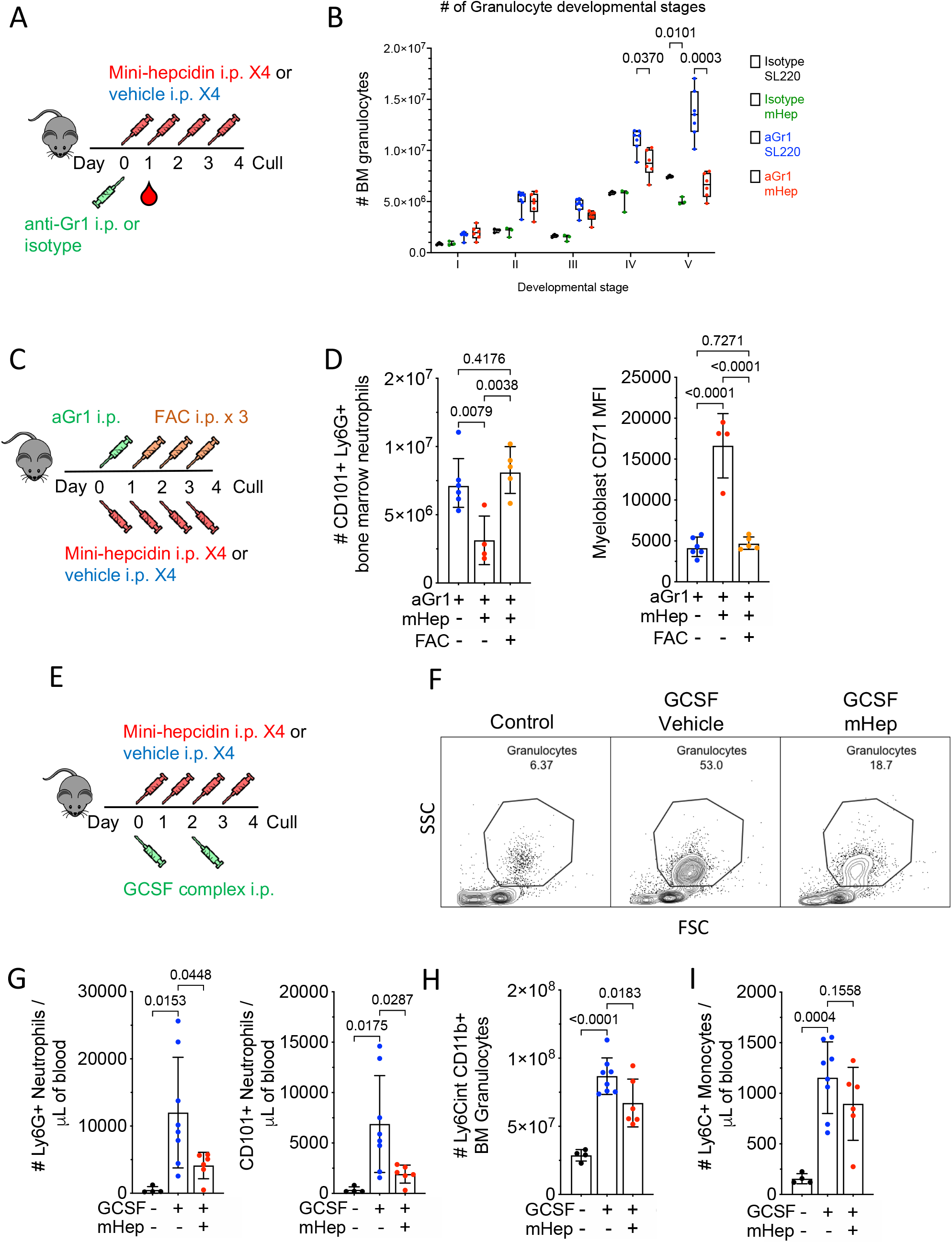
Control of enhanced myelopoiesis by hepcidin driven iron deficiency. A) Experimental scheme for effect of minihepcidin treatment on recovery after neutrophil depletion B) Bone marrow granulocyte differentiation trajectory in mice treated as in Fig. 2A, representative to two independent experiments. Two way ANOVA. Mean, quartiles and range. C) Experimental scheme for stydying the effect of mHep-induced hypoferremia and its amelioration with ferric ammonium citrate (FAC) on recovery from aGr1-mediated neutrophil depletion. All mice received anti-Gr1. D) Number of mature bone marrow CD101+ neutrophils and myeloblast CD71 expression from mice treated as in Fig 2C. One way ANOVA. Mean ± SD. E) Experimental scheme for studying the effect of mHep on GCSF complex driven granulocyte expansion F) Example flow cytometry scatter plot indicating frequency of neutrophils in the blood from experiment outlined in 2E. G) Number of blood neutrophils in the blood from experiment outlined in 2E. One way ANOVA. Mean ± SD. H) Total bone marrow granulocytes and CXCR4+ C-kit+ granulocyte precursors in the bone marrow from experiment outlined in 2E. One way ANOVA. Mean ± SD. I) Number of blood monocytes in the blood from experiment outlined in 2E. One way ANOVA. Mean ± SD.

**Figure 3.**
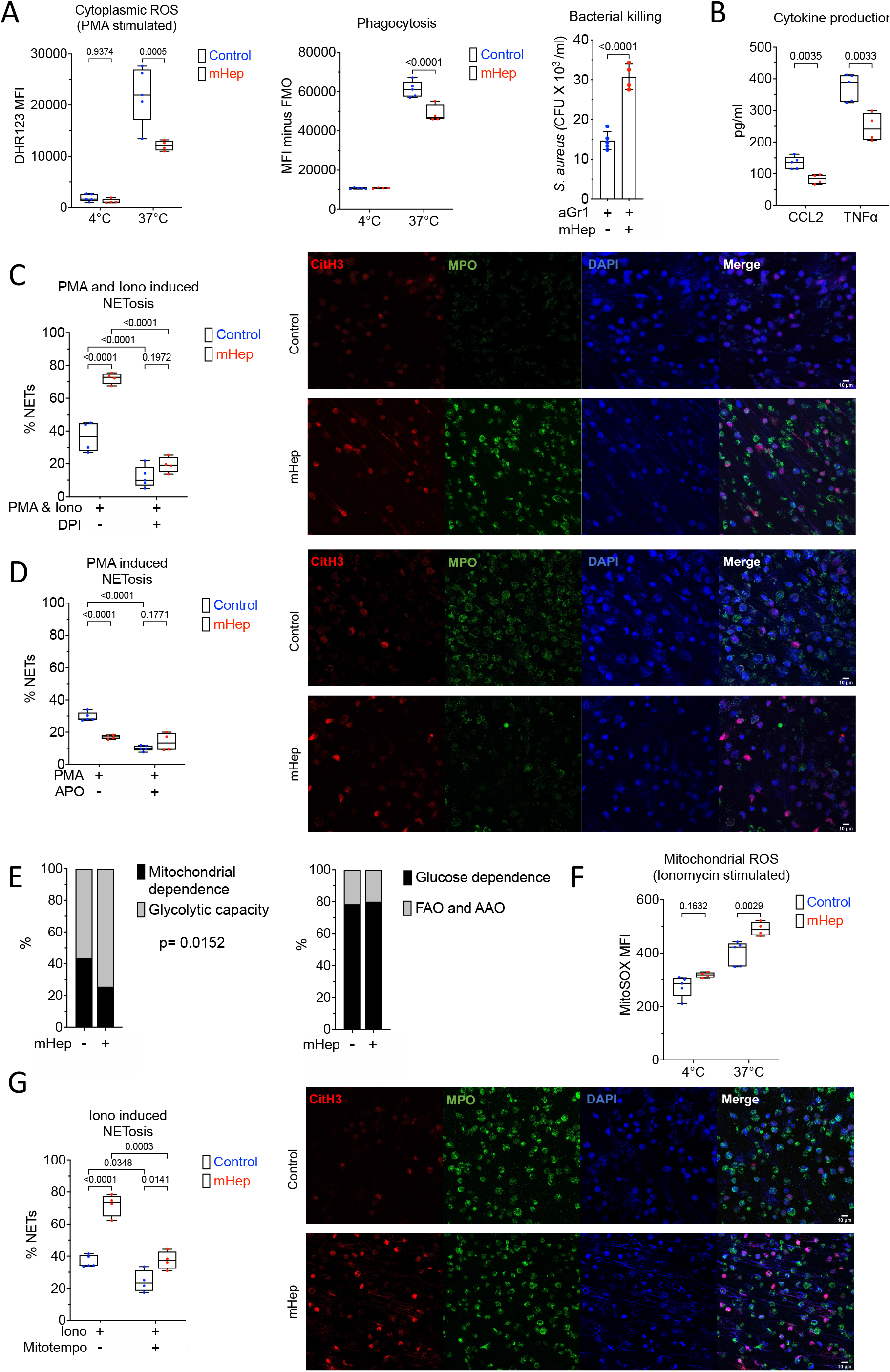
Functional alteration of neutrophils in serum iron deficiency. A) Phenotyping *ex vivo* of isolated bone marrow neutrophils (obtained from mice treated as in Figure 2A). DHR123 fluorescence as a reporter for cytoplasmic ROS in PMA stimulated cells. Phagocytosis of GFP labelled E.coli. Two way ANOVA. Mean, quartiles and range. Bacterial killing of *S. aureus*, resolved as CFU of S. aureus recovered after co-culture with neutrophils. Unpaired T-test. Mean ± SD. B) Supernatant CCL2 and TNF levels produced by zymosan stimulated neutrophils measured by ELISA. Unpaired T-test. Mean, quartiles and range. C) NETosis was evaluated in isolated bone marrow neutrophils after stimulation ex vivo with PMA and ionomycin ± diphenyleneiodonium chloride (DPI). A minimum of 250 cells from two replicates per sample counted. Two way ANOVA. Mean, quartiles and range. Representative microscopy images taken at 20X magnification. D) NETosis was evaluated in isolated bone marrow neutrophils after stimulation ex vivo with PMA ± apocynin (APO). A minimum of 250 cells from two replicates per sample counted. Two way ANOVA.. Mean, quartiles and range. Representative microscopy images taken at 20X magnification. E) SCENITH analysis of Ly6G+ bone marrow neutrophils from experiment described in 1A, analysing proportional shifts in OPP incorporation after ex vivo treatment with metabolic inhibitors as detailed in methods. Two way ANOVA with matching for sample. Reported p-value is the effect of mHep treatment. Mean. F) MitoSOX fluorescence as a reporter for mitochondrial ROS in ionomycin stimulated isolated bone marrow neutrophils. Two way ANOVA. Mean, quartiles and range. G) NETosis was evaluated in isolated bone marrow neutrophils after stimulation ex vivo with ionomycin (iono) ± mitoTEMPO. A minimum of 250 cells from two replicates per sample counted. Two-way ANOVA. Mean, quartiles and range. Representative microscopy images taken at 20X magnification.

### Counteracting inflammatory hepcidin induction improves neutrophil production

Low plasma iron and raised hepcidin are commonly observed in the context of inflammation (Weiss et al., 2019). Having demonstrated iron-dependent granulopoietic suppression under non-inflammatory conditions, we challenged mice after 4 days of mHep treatment with lipopolysaccharide (LPS) to interrogate how intensified hypoferremia alters the neutrophil response in the context of inflammation (4A). Hypoferremia reduced inflammatory accumulation of neutrophils in the spleen and peritoneal cavity (4B); however, no difference in the number of Ly6G+ neutrophils remaining in the BM was observed (S4A). This is possibly because inflammatory mobilisation out of the BM dominated over production in these conditions.

We next considered whether endogenous hepcidin would suppress granulopoiesis during inflammation. Hepatic hepcidin transcription is induced during inflammation by IL-6 via a signalling axis that also requires sustained tonic BMP6/SMAD signalling (Wang and Babitt, 2019). After depleting neutrophils using anti-GR1 to enhance granulopoietic demand, we administered mice with LPS as a hepcidin-stimulatory inflammatory signal, with or without anti-BMP6 antibody (4C). As previously shown (Petzer et al., 2020), BMP6 neutralisation suppressed the endogenous hepcidin response to inflammation and increased plasma iron at 24hrs post-LPS injection (4D). The LPS dose used here previously caused transient hepcidin-dependent hypoferremia at 6hrs post-injection that was resolved by 24hrs (Armitage et al., 2016). Accordingly, in the present experiment, although serum iron in animals treated with LPS alone was similar to that of controls at 24hrs, these mice had likely experienced transient inflammatory hypoferremia (4D). Although anti-BMP6 increased serum iron, it did not alter other aspects of systemic LPS-mediated inflammation, as determined by weight loss (S4B), liver *Saa1* and *Fga* mRNA expression (S4C) and serum cytokine levels (S4D).

Focusing on granulopoiesis, anti-BMP6 increased numbers of mature Ly6G+ neutrophils and CD115-ve c-kit+ neutrophil progenitors in the bone marrow of inflamed mice (4E and 4F). Furthermore, CD71 expression by both Ly6G+ cells and progenitors was decreased in anti-BMP6 treated mice, indicating that improved iron acquisition by granulopoiesis during inflammation was associated with increased production of neutrophils (4E and 4F).

## Discussion

Systemic hypoferremia is a feature of the acute phase response of human infections (Armitage et al., 2014; Darton et al., 2015; Shah et al., 2020; Williams et al., 2019) and likely protects against extracellular siderophilic infections. However, beyond the well-characterised anaemia of inflammation (Weiss et al., 2019), further consequences of hypoferremia for host physiology are poorly defined. We show that granulopoiesis is particularly sensitive to iron availability, commensurate with the relatively high production of neutrophils containing a relatively iron-rich proteome.

In contrast to granulopoiesis, steady-state and GCSF-stimulated monocyte production appeared resistant to hypoferremia, with evidence of increased numbers of bone marrow monocyte and their progenitors in resting mice. Monocyte cytokine production *ex vivo* was also preserved. This is a striking result as, like neutrophils, monocytes are an innate cell type exhibiting rapid turnover *in vivo* (Evrard et al., 2018; Yona et al., 2013), supporting our hypothesis that iron may be particularly important to neutrophils beyond the requirement to support the production of a cell type with a short half-life. Notably the ‘iron resistance’ of the monocyte lineage is supported by observations of normal NO_2_ production by LPS stimulated monocytes from iron deficient individuals (Paino et al., 2009).

Our results suggest that neutrophil progenitors have higher baseline iron requirements than monocyte progenitors, and that iron restriction does not perturb monocyte progenitor cell-cycle profiles. Functionally, hypoferremia could modulate the overall profile of innate immunity at baseline and during stress myelopoiesis by controlling the ratio of monocytes to neutrophils. Interestingly, in addition to these systemic effects, local skin derived hepcidin also influences innate immunity by facilitating mature neutrophil chemotaxis (Malerba et al., 2020).

We found that neutrophils produced during hypoferremia exhibited reduced ability to phagocytose fluorescent *Escherichia coli*, produce cytokines and kill *Staphylococcus aureus*. We also observed suppression of NOX-dependent NETosis, linking systemic nutrient availability to NETosis and building on previous reports of inhibited NOX-dependent ROS production during iron deficiency (Ahluwalia et al., 2004; Kurtoglu et al., 2003; Paino et al., 2009). Although non-physiological iron chelation has previously been linked to NETosis with conflicting results (Kono et al., 2016; Völlger et al., 2016), we observe that physiological iron restriction greatly enhanced mitoROS production and mitoROS dependent NETosis. The importance of different NETosis pathways in response to different stimuli is complex and an ongoing area of study (Kenny et al., 2017). However, notably, mitoROS-dependent NETosis may play a dominant role in immune complex driven NETosis during Lupus-like autoimmunity (Lood et al., 2016), where NETs play a key role in driving pathology (Gupta and Kaplan, 2021). Such patients frequently present with raised hepcidin and low serum iron (Kunireddy et al., 2018). Our results prompt the question of whether prolonged iron restriction due to raised hepcidin under chronic inflammatory conditions contributes to neutrophil dysfunction, NETosis and propagation of immunopathology in autoimmunity.

Consistent with our observations in T cells (Frost et al., 2021), we show that iron restriction changes haematopoietic cell metabolism *in vivo*, suppressing oxidative mitochondrial metabolism in bone marrow granulocytes. Disrupted mitochondrial metabolism may be linked to increased mitoROS production. Whilst mature neutrophils are extremely reliant on glycolysis for ATP production (Borregaard and Herlin, 1982), oxidative mitochondrial metabolism – which is a highly iron dependent process - has been suggested to be important for neutrophil differentiation and development in the bone marrow (Riffelmacher et al., 2017). The impaired metabolism, differentiation and function of iron deficient neutrophils in our study is consistent with this. Further studies will be required to elucidate how iron mechanistically supports neutrophil differentiation, biosynthesis of effector proteins, metabolism and progenitor proliferation.

Blocking hepcidin and increasing serum iron in a mouse model of acute inflammatory hypoferremia increased the number of bone marrow granulocytes without altering markers of inflammation. We hypothesise that therapeutic hepcidin suppression could be leveraged to control neutrophil numbers and functionality during infection with pathogens insensitive to hepcidin-mediated hypoferremia (Stefanova et al., 2017) or during autoimmunity where raised hepcidin might contribute to neutrophil dysfunction and pro-inflammatory NETosis.

Relatively little is known regarding how changes in systemic nutrient availability, including hypoferremia, modulates innate immune cell function *in vivo*. Our results highlight that neutrophil production is a particularly iron-dependent aspect of haematopoiesis, sensitive to rapid physiological changes in serum iron concentration. We propose that hepcidin-mediated hypoferremia is a novel immune axis regulating myelopoiesis, specifically negatively regulating granulopoiesis, resulting in fewer neutrophils with altered effector functions. In contrast, monopoiesis is unscathed by hypoferremia. These findings have implications for understanding inflammatory pathology and for the clinical use of granulopoiesis-mobilising agents in patients at risk of infection and iron deficiency.

## Acknowledgements

The authors thank the staff of the University of Oxford Department of Biomedical Services for animal husbandry, Yi-Ling Chen for assistance running the Luminex assay; Goran Mohammad and Samira Lakhal-Littleton for assistance with serum iron measurements.

This work and J.N.F., A.E.P., A.E.A. and H.D. was supported by the UK Medical Research Council (MRC Human Immunology Unit core funding to HD, award no. MC_UU_12010/10). S.K.W. was supported by the Wellcome Trust Infection, Immunology & Translational Medicine doctoral programme (grant no. 108869/Z/15/Z). M.R.T. was supported by the Clarendon Fund and the Corpus Christi College A. E. Haigh graduate scholarship.

A.C. was supported by an Oxford-BMS Fellowship. B.G. is supported by grants from the Deutsche Forschungsgemeinschaft (GA2075/5-1, GA2075/6.1). A.J.C. is supported by the Wellcome Trust (211072/Z/18/Z). This work was supported by the Novo Nordisk Foundation (Tripartite Immunometabolism Consortium to L.W. and MeRIAD Consortium to I.A.U.), Chinese Science Council (to Z.A.), and the Wellcome Trust (Investigator Award 209422/Z/17/Z to I.A.U.).

## Methods

### Resource availability Lead contact

Further information and requests for resources and reagents should be directed to and will be fulfilled by the Lead Contact, Hal Drakesmith

### Materials availability

This project did not generate new unique reagents

### Data and code availability

This project did not generate any new code or novel datasets

### Experimental models and subject details

#### Mice

Unless otherwise stated, animal procedures were performed under the authority of UK Home Office project and personal licenses in accordance with the Animals (Scientific Procedures) Act 1986, and were approved by the University of Oxford ethical review committee. Mice were housed in individually ventilated cages and fed ad-libitum with a standard diet containing 188ppm iron (SDS Dietex Services, diet 801161). Mice were euthanised in increasing CO_2_ concentrations. All mice were sex matched and age matched (to within 2-weeks) within individual experiment. WT C57BL/6JOlaHsd mice were ordered from Envigo. Mice were used between 6-10 weeks of age when initiating each experiment. Female mice were used for all experiments, aside from experiment described in figure 4C where male mice were used because of their lower liver iron stores and more pronounced acute phase response. Within each experiment mice were randomly allocated to treatment groups such that an equal number of mice in each cage received each treatment.

**Figure 4.**
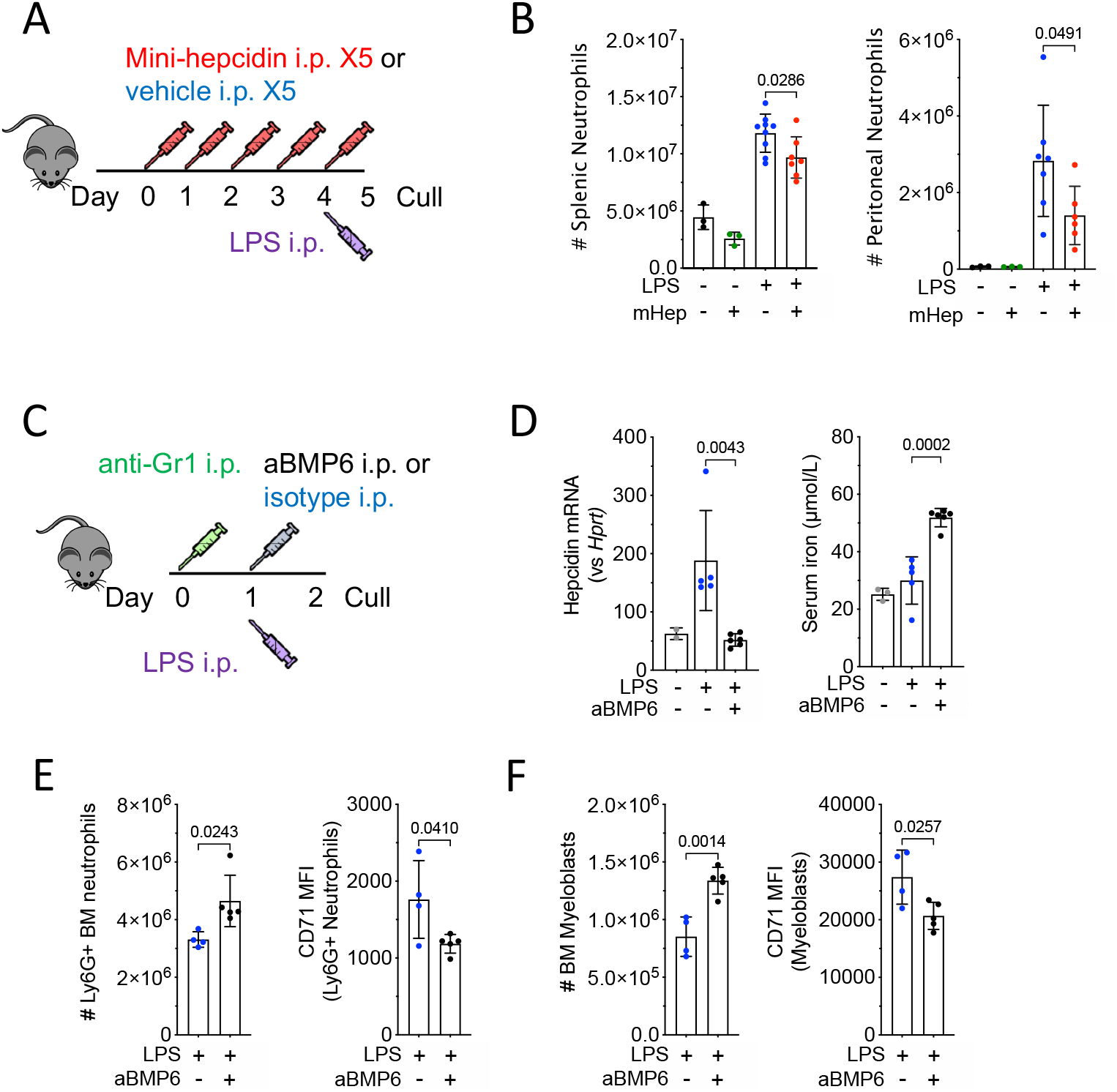
Modulation of hepcidin during inflammation alters neutrophil production. A) Experimental scheme for investigation of effect of sustained hypoferremia on neutrophil response to sterile inflammatory challenge. B) Number of mature Ly6G+ neutrophils in the spleen and peritoneal cavity in mice treated as in 4A. t-test. Mean ± SD. C) Experimental scheme for investigation of effect of endogenous hepcidin response on granulopoiesis. D) Liver hepcidin expression (Mann-Whitney test) and serum iron from mice treated as in 4C. Mean ± SD. Non LPS treated control group was treated with aGr1 24hrs prior to culling to reflect the systemic iron environment prior to LPS injection. Representative of two independent experiments E) Number of bone marrow Ly6G+ neutrophils and CD71 MFI on Ly6G+ neutrophils in mice treated as in 4C. t-test. Mean ± SD. Representative of three independent experiments. F) Number of BM c-kit+ CD115-Ly6G-myeloblasts and CD71 MFI on c-kit+ myeloblasts in mice treated as in 4C. t-test. Mean ± SD. Representative of three independent experiments.

#### Injected substances

Minihepcidin: An appropriate mass of mini-hepcidin PR73 (da-TH-Dpa-bhPro-RCR-bhPhe-Ahx-Ida(Hexadecylamine)-NH2) was dissolved in 80% ethanol and then mixed with 60mg of Purebright SL220/ Sunbright DSPE-020CN (NOF). The control solution was Purebright SL220 dissolved in ethanol. The ethanol was evaporated off using a vacuum chamber warmed to 50°C. The resultant gel was stored up to 24 hours at 4°C and re-dissolved in an appropriate volume of water to give a mHep concentration of 1mM: 100nMoles of mini-hepcidin in 100μl of water was injected per mouse per dose.

Ferric ammonium citrate (FAC) (F5879 Sigma Aldrich) was dissolved to 3mg/ml in PBS. 150μl was injected intraperitoneally into mice. Anti-Gr1-mediated neutrophil depletion: 10μg (female mice) or 20μg (male mice) of anti-Gr1/mouse (Ultra-LEAF, clone RB6-8C5, Biolegend, 108436) diluted in sterile low endotoxin PBS was injected intraperitoneally into mice. *E*.*coli* LPS-055:B5 (Sigma, L2880) was injected at a dose of 1mg/kg body weight diluted in sterile low endotoxin PBS (Armitage et al., 2016) and injected intraperitoneally into mice. Anti-BMP6 and isotype control were injected at a dose of 10mg/kg body weight diluted in sterile low endotoxin PBS and injected intraperitoneally into mice.G-CSF complexes were prepared as described in (Rubinstein et al., 2013) using human G-CSF (Filgrastim) and anti-human G-CSF (Clone BVD11-37G10, Southern biotech, Cat. No. 10128-0). 7.5μg of anti-G-CSF and 1.5μg of G-CSF were complexed, diluted in sterile low endotoxin PBS and injected intraperitoneally. 7.5μg of aG-CSF alone was used as the control injection to control for trace endotoxin contamination of the antibody and direct effects of the antibody.

#### Flow cytometry staining

Single cell suspensions of spleen were made by mechanical dissociation through a 40μm cell strainer. Red cells were lysed in spleen suspensions using Tris ammonium chloride (ACT) red cell lysis buffer (2.06g/L Tris base and 7.47g/L NH_4_Cl, 1L H_2_0, adjusted to pH 7.2). Bone marrow was flushed though a 70μm filter to make a single cell suspension. Peritoneal lavage was obtained by flushing the peritoneal cavity with a 23g needle and syringe filled with 8ml of cold PBS.

For analysis of murine peripheral blood leukocytes, 50-100μl of whole blood collected by tail bleed into a BD microtainer EDTA tube was mixed with 1ml of ACT red cell lysis buffer and incubated at room temperature for 20 minutes. The blood solution was spun down at 400g for 5 minutes, supernatant removed, and the leukocyte pellet transferred to a round bottom 96 well plate for flow cytometric staining.

Tissue cell suspensions or *in vitro* activated cells at appropriate time points after activation cells were transferred to a 96 well round bottom plate, spun down and washed with 200μl of PBS. Cells were stained with appropriate concentrations of FC receptor block, fluorophore conjugated antibodies and a fixable live dead dye in 40μl of PBS for 20 minutes at 4°C in the dark. Cells were washed and ran directly on an Invitrogen Attune or BD LSR Fortessa. For some experiments, cells were fixed for 10 minutes in 100μl of 4% paraformaldehyde at 4°C in the dark before washing and resuspension for analysis or resuspended in saponin based perm buffer to stain intracellular antigens. Intracellular Ly6G staining was used 24hrs after aGr1 treatment to accurately resolve neutrophil depletion despite masking of surface epitopes (Gael Boivin 2020).

DAPI (1μg/ml) was added prior to running, to fixed and permeabilised cells, for DNA content staining.

#### Cell culture

##### All cells were cultured at 37°C, 20% O_2_ and 5% CO_2_

For monocyte intracellular cytokine production 2 million whole splenocytes (in a round bottom 96 well plate) were activated for 3 hours in the presence of Brefeldin A (5μg/ml) with/without LPS (5ng/ml) in iron free medium. Iron free medium: RPMI, glutamine (2mM), penicillin (100U/ml), streptomycin (0.1mg/ml) and Beta-mercaptoethanol (55μM), to preserve *in vivo* iron status FCS was replaced with 10% v/v Pannexin NTS serum substitute (P04-95080; Pan Biotech, custom order). Splenocytes were then washed and stained for surface epitopes and intracellular cytokines as described in flow cytometry.

#### Neutrophil isolation

Neutrophils were negatively-isolated from whole bone marrow of minihepcidin treated and control treated animals as per the manufacturer’s instructions (Neutrophil Isolation Kit, 130-097-658, miltenyi biotec).

#### Flow cytometric ROS measurement

To identify DHR123 staining of predominantly cytoplasmic ROS, isolated neutrophils were incubated with 2.5 μg/mL (7 μM) Dihydrorhodamine 123 (DHR123) (ThermoFisher Scientific) in complete RPMI 1640 medium, and stimulated by 50nM Phorbol 12-Myristate 13-Actetate (PMA) (Sigma-Aldrich) for 20 minutes at 37°C. Cells were subsequently washed with PBS and the fluorescence intensities of each subset/cells were measured by flow cytometry. 4°C incubation was used as controls.

Mitochondrial ROS (mitoROS) was measured using mitoSOX™ red (ThermoFisher Scientific). isolated neutrophils in complete RPMI 1640 medium were incubated with 5μM mitoSOX red for 20 minutes at 37°C in the presence of 10 μM ionomycin. Cells were subsequently washed with PBS and the fluorescence intensity as the indicator of mitoROS generation were measured by FACS. 4°C incubation was used as controls.

#### Bacterial killing assay

Bacterial killing assay was performed with S. aureus (NCTC 6571), which was used at a MOI of 10. For the bacterial killing assay, isolated neutrophils were interacted with S. aureus for **2 hours** at 37 °C in 5% CO^2^ tissue culture incubator and then lysed in 1% triton buffer. The lysates were then plated on agar plates. Bacterial culture plates were incubated at 37 °C overnight, and the colony number on each plate was counted the following morning as an absolute CFU count.

#### Measurement of cytokine production

For the cytokine array, 2×10^6^ isolated neutrophils seeded in 2ml RPMI were stimulated with 50ug/ml Zymosan or DMSO vehicle for two hours. After incubation, Mouse CCL2 DuoSet Elisa (R&D Systems, catalog# DY479-05), Mouse TNF-alpha DuoSet Elisa (R&D Systems, catalog MTA00B) were used to detect the levels of CCL2 and TNF-alpha, respectively, in the supernatants of stimulated isolated neutrophils, following the manufacturer’s instruction. For readout, signals were detected by chemical luminescence and subsequently measured with Microplate Reader.

#### NETosis

To induce NETosis, isolated neutrophils were seeded into 8 well lab-Tek II chamber slide (VWR international) coated with 2% poly-lysine (Sigma-Aldrich) at a volume of 100 μL at the density of 1×10^6^/mL. Neutrophils were stimulated with 10 μM ionomycin and 10 μM PMA together or individually, overnight at 37°C in 5% CO^2^ tissue culture incubator and were subsequently fixed with 4% paraformaldehyde (Sigma-Aldrich) in DPBS for 30 minutes at RT. To investigate the effect of ROS on NETosis, different ROS inhibitors were used. 25 μM diphenyleneiodonium chloride (DPI) was used to block total ROS generation from ionomycin and PMA, 100 μM apocynin was used to block ROS generation via NOX2 complex by PMA and 5 μM mitoTEMPO (Cayman Chemical) was used to inhibit mitoROS production. Afterwards, cells were washed three times with DPBS, and incubated with blocking buffer (2% BSA in TBST) for 20minutes. Following blocking, primary antibodies: rabbit anti-citrullinated histone 4 (Abcam, ab5103) and mouse anti-mouse MPO (Hycult, HM1051BT) at 1:100 dilution in blocking buffer, were added and incubated for 2 hours at RT or overnight at 4°C. Cells were washed with DPBS before adding secondary antibodies: mouse anti rabbit DyLight 647 conjugated secondary antibody (ThermoFisher Scientific) and rabbit anti mouse IgG secondary antibody conjugated with Alexa fluor 488 (ThermoFisher Scientific) at 1:300 dilution in blocking buffer for an 1h incubation at RT in the dark. Subsequently, cells were washed with DPBS. Before imaging, the chambers were removed from the slides and cells were covered with ProLong™ gold antifade mounting media with DAPI (ThermoFisher Scientific). Images were obtained by Zeiss LSM 980 confocal microscope. Neutrophils with a clear formation of fibres, together with a diffuse nucleus stained by DAPI and colocalization with MPO and citrullinated-histone 3, were counted as neutrophils under NETosis. For each condition, 100-200 cells of each sample have been counted from three different fields of two independent experiments. ImageJ was used for image analysis and presentation.

#### SCENITH analysis

SCENITH is a flow cytometry approach in which translation rate is quantified by measuring puromycin incorporation into protein (Argüello et al., 2020; Lopes et al., 2021). Translation is highly ATP dependent. Therefore, changes in the rate of translation after the addition of different metabolic inhibitors can be used to estimate cellular glucose dependence, mitochondrial dependence, glycolytic capacity and fatty acid / amino acid oxidation capacity. The SCENITH protocol was adapted from the original method designed by Arguello et al. (Argüello et al., 2020) for use with the Invitrogen Click-iT Plus OPP protein synthesis assay kit (C10456, Invitrogen). To prevent cells from iron loading *ex vivo* during short term culture, all steps were conducted in iron free media. Bone marrow cells were plated at 3×10^6^ cells per well in a 96 well plate. Matched samples were treated in parallel with 100 μL of iron free media at 37°C, 5% CO_2_ with 5 different conditions: media alone, 2-DG (100mM), oligomycin (1 μM), 2-DG (100mM) + oligomycin (1 μM), and harringtonin (2 μg/mL). 30 minutes into inhibitor treatment, 100 μL of O-propargyl-puromycin (OPP) was added on top for a final OPP dilution of 1:1000 and incubated for a further 30 minutes at 37°C, 5% CO_2_. Cells were then washed, surfaced stained, fixed and permeabilised as described in flow cytometry. Following permeabilization, intracellular OPP was labelled according to the manufacturer’s instructions in the the Invitrogen Click-iT Plus OPP protein synthesis assay kit (C10456, Invitrogen). Analysis was conducted as described by Arguello et al.

#### Blood measurements

Full murine red blood cell indices and platelet counts were performed on 100μl of cardiac whole blood taken into a BD EDTA microtainer (Beckton Dickinson) and measured on a Sysmex KN-21 blood analyser.

For serum analysis of murine samples up to 400μl of blood obtained by cardiac puncture was placed in a BD microtainer SST tube (Beckton Dickinson). Serum was obtained by spinning the clotted blood sample was spun at 8,000g for 5 minutes and stored at -80°C. Iron measurements were determined using an ABX Pentra instrument. Serum cytokine levels were measured by Luminex (IL-5, GM-CSF and G-CSF, life technologies) or Legendplex (M1 macrophage panel, with added G-CSF capture beads,740848, Biolegend) in accordance with the manufacturer’s instructions.

#### Mathematic modelling

Iron counts per cell were calculated using the method described in Teh et al. (Teh et al., 2021). Human iron interacting proteins were identified in human immune cell proteomes (Derived from (Rieckmann et al., 2017)) by cross-referencing against the list of human iron interacting proteins identified by Andreini et al. (Andreini et al., 2018)). Hemoglobin proteins (HBA1, HBB, HBD, HBG1, HBG2, HBM, HBQ1, and HBZ) were removed as these proteins should not be expressed in immune cells and their presence is likely indicative of red cell contamination. Protein copy number values for each iron interacting protein species was multiplied against the protein:iron atom stoichiometry values previously curated in Teh et al. (Teh et al., 2021) to give the number of iron atoms per protein species. Total cellular iron counts were calculated as the sum of iron atoms attributed across all detected iron interacting protein species.

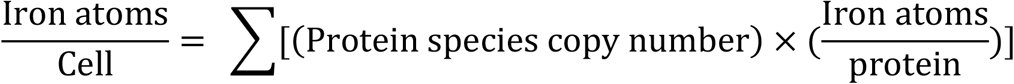

We assigned iron-interacting protein species to pathways using pre-established GO and Reactome gene sets. OXPHOS: GOBP_AEROBIC_RESPIRATION; DNA replication: GOBP_DNA_REPLICATION; iron pathways: pooled GOBP_CELLULAR_IRON_ION_HOMEOSTASIS and GOBP_IRON_SULFUR_CLUSTER_ASSEMBLY; demethylation: GOBP_DEMETHYLATION; Fatty Acid metabolism: GOBP_FATTY_ACID_METABOLIC_PROCESS; Granule proteins: REACTOME_NEUTROPHIL_DEGRANULATION. Where overlaps in gene sets were present, we assigned pathways according to best fit given the current literature (see scripts for specific designations). Atoms attributed to protein species were in turn assigned to each protein’s corresponding pathway.

---

**Figure S1.**
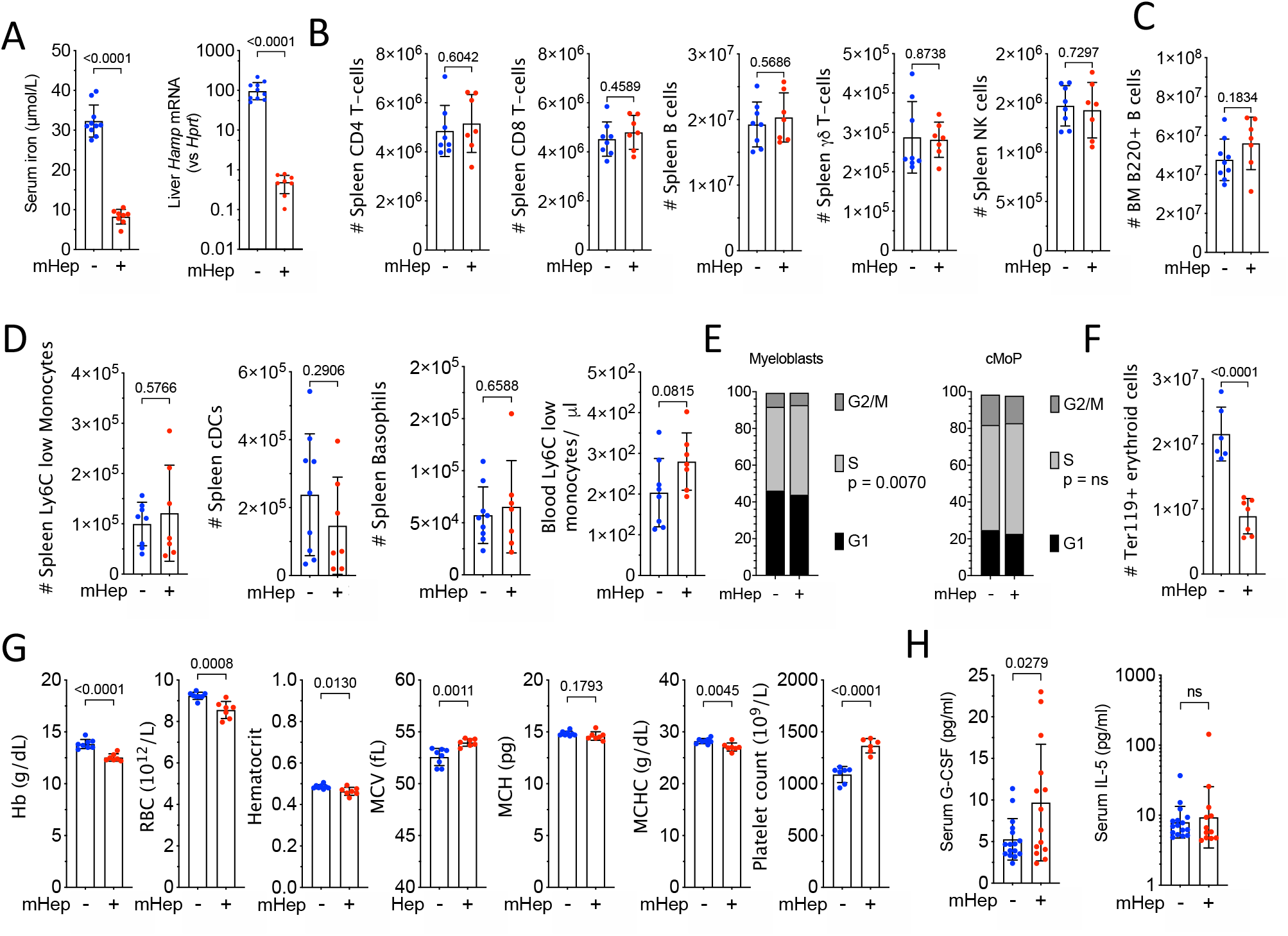
A) Serum iron and liver hepcidin mRNA in experiment described in Fig 1C, Representative of two independent experiments. Unpaired T-test. Serum iron, mean ± SD. Hepcidin mRNA, geometric mean ± geometric SD. B) Number of splenic lymphocyte in experiment described in Fig 1C. Representative of two independent experiments. Unpaired T-test. Mean ± SD. C) Number of B220+ B cells in the bone marrow in experiment described in Fig 1C. Unpaired T-test. Mean ± SD. D) Number of specified splenic and blood myeloid cells in experiment described in Fig 1C. Representative of two independent experiments. Unpaired T-test. Mean ± SD. E) Cell cycle distribution of myeloblast and cMoP in experiment described in Fig 1C. Representative of two independent experiments. Two way ANOVA, mean proportions of the whole. F) Number of Ter119+ erythroid cells in the bone marrow in experiment described in Fig 1C. Unpaired T-test. Mean ± SD. G) Red cell indices and platelet count were measured by sysmex in experiment described in Fig 1C. Unpaired T-test. Mean ± SD. H) Serum G-CSF and IL-5 were measured by Luminex in experiment described in Fig 1C. 2-way ANOVA blocking on experimental repeat and reporting p-value treatment effect, combination of two independent experiments. G-CSF, mean ± SD. IL-5, geometric mean ± geometric SD.

**Figure S2.**
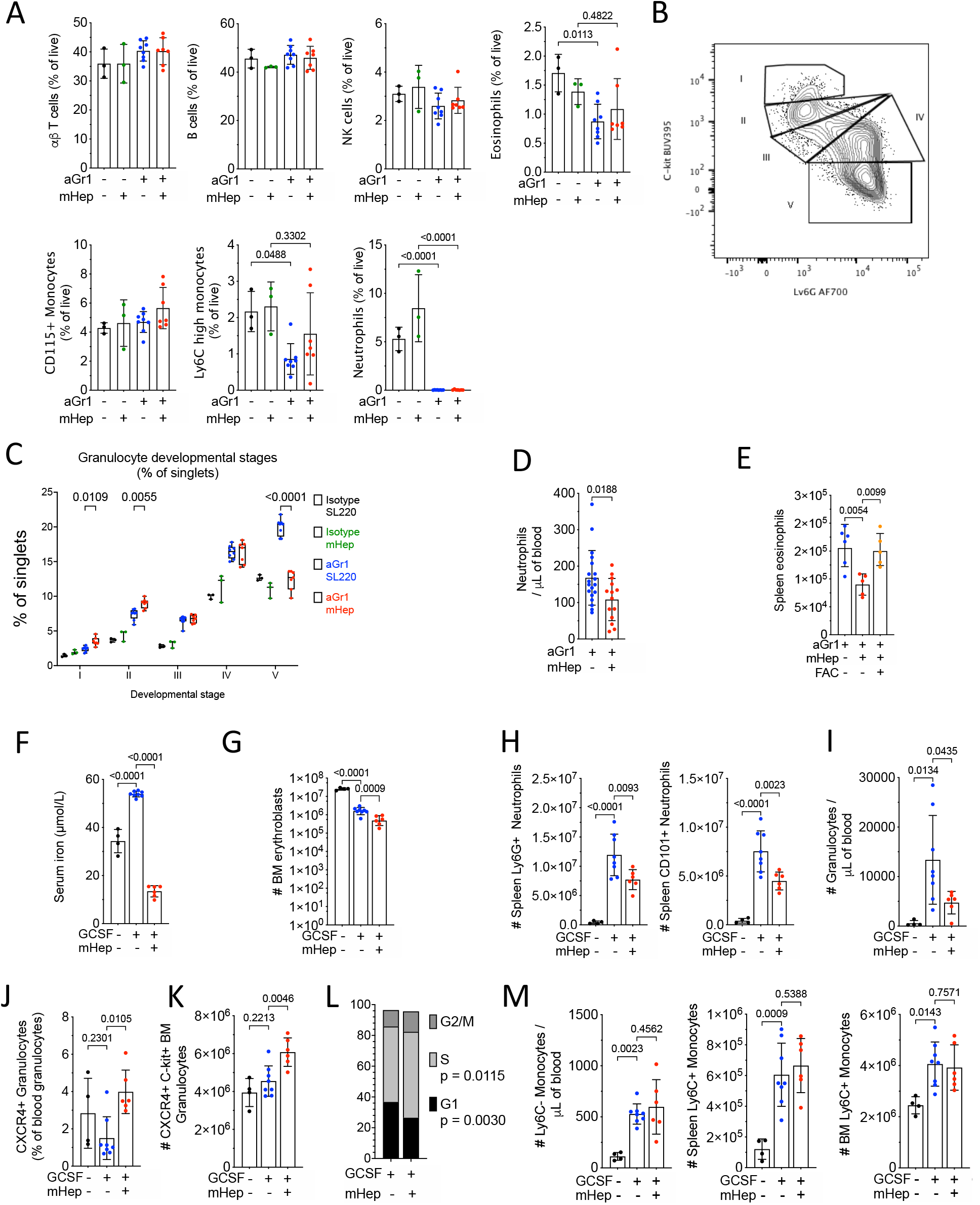
A) Frequency of blood leukocyte subsets as a % of live events 24hrs after 10μg of aGr1 or isotype control as part of experiment outlined in Fig 2A. Two way ANOVA. Mean ± SD B) Gating scheme to delineate stages of granulocyte differentiation (gated on Ly6C+, CD11b+, CD115-cells) in Fig 2B. From stage I (myeloblasts) to stage IV (mature neutrophils). C) Bone marrow granulocyte differentiation trajectory in mice treated as in Fig S2A, representative to two independent experiments. Two way ANOVA. Mean, quartiles and range. D) Blood neutrophil numbers in mice treated as in scheme 2A (combined from three independent experiments). Unpaired T-test. Mean ± SD E) Spleen Eosinophil numbers in mice treated as in scheme 2C. One way ANOVA. Mean ± SD F) Serum iron concentrations in experiment scheme 2E. One way ANOVA. Mean ± SD G) Bone marrow CD71+ TER119+ erythroblast numbers in scheme 2E. One way ANOVA. Geo Mean ± GeoSD H) Spleen Neutrophil numbers in experiment scheme 2E. One way ANOVA. Mean ± SD I) Blood Ly6C int CD11b+ granulocyte numbers in experiment scheme 2E. One way ANOVA. Mean ± SD J) % of granulocytes showing an immature CXCR4+ phenotype in experiment scheme 2E. One way ANOVA. Mean ± SD K) Number of Bone marrow CXCR4+ c-kit+ neutrophil progenitors in experiment scheme 2E. One way ANOVA. Mean ± SD L) Cell cycle distribution of CXCR4+ c-kit+ neutrophil progenitors in experiment scheme 2E. Two way ANOVA. Mean proportion of whole. M) Monocyte numbers in blood, spleen and bone marrow in experiment scheme 2E. One way ANOVA. Mean ± SD

**Figure S3.**
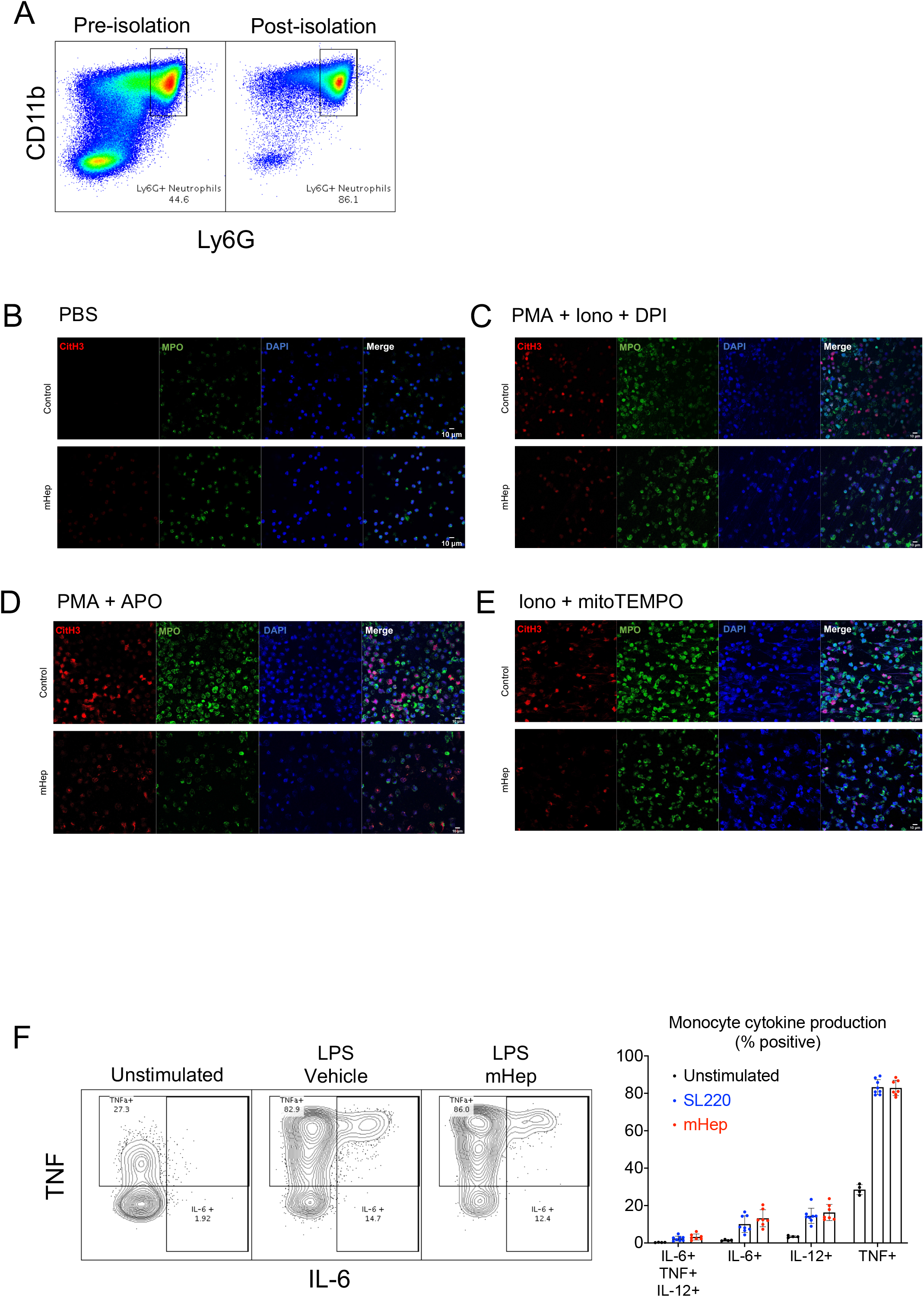
Functionality of myeloid cells in serum iron deficiency. A) Enrichment purity of bone marrow Ly6G+ neutrophils (obtained from mice treated as in experiment scheme Fig 2A) Representative microscopy images of NETosis taken at 20X for the following conditions B) PBS treatment C) PMA, Ionomycin and DPI D) PMA and apocyanin (APO) E) Ionomycin and mitoTEMPO B) Intracellular cytokine staining within Ly6C+ CD11b+ monocytes after *ex vivo* LPS stimulation of whole splenocytes from mice treated as in experiment scheme 1C. One way ANOVA. Mean ± SD

**Figure S4.**
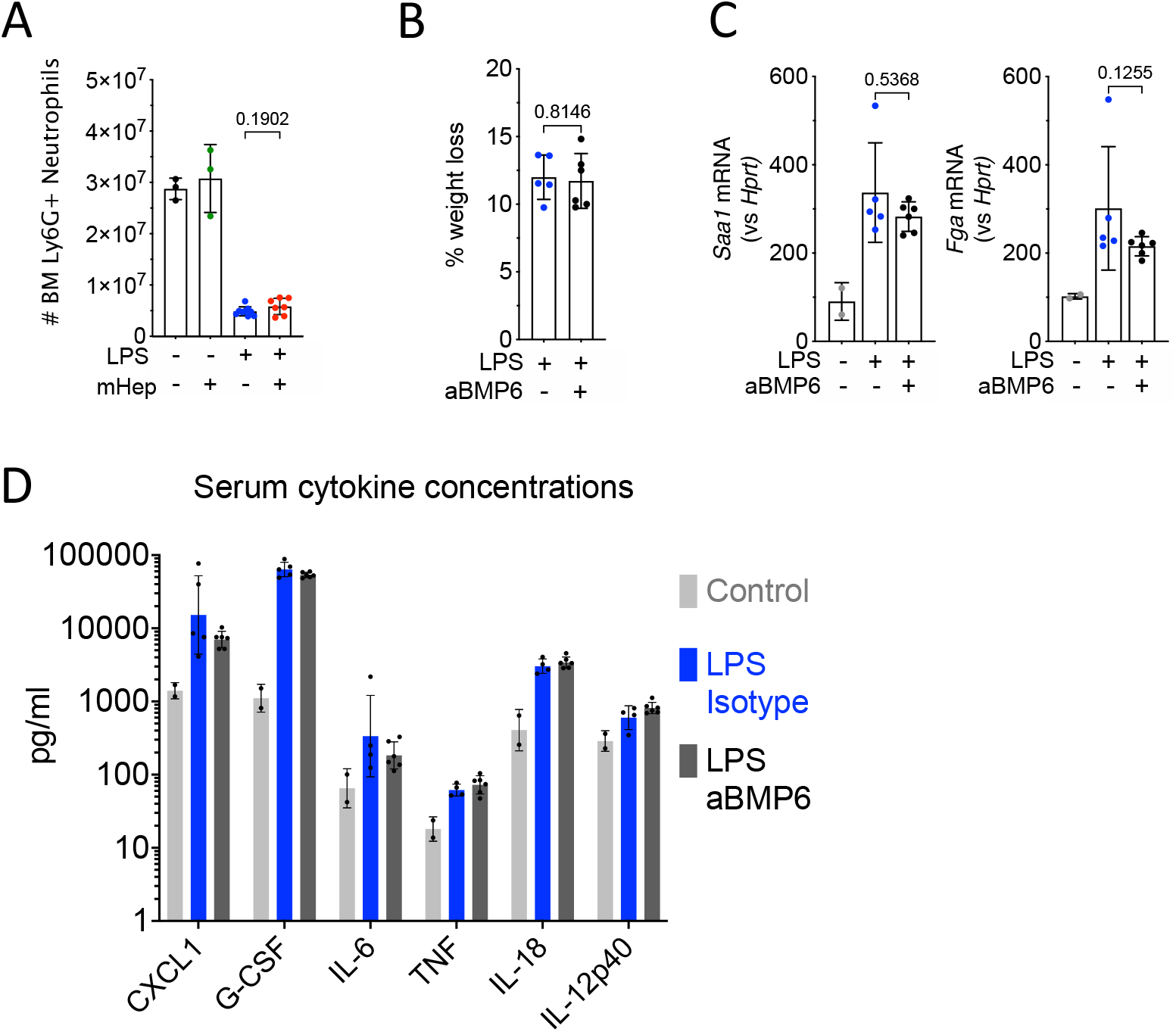
Counteracting inflammatory hepcidin induction improves neutrophil production. A) BM neutrophil numbers from mice treated as in 4A. t-test. Mean ± SD. B) LPS induced weight loss from mice treated as in 4C. t-test. Mean ± SD. Representative of three independent experiments C) Liver Saa1 and Fga expression (Mann-Whitney test) from mice treated as in 4C. Mean ± SD. Non LPS treated control group was treated with aGr1 24hrs prior to culling to reflect the
systemic iron environment prior to LPS injection. Representative of two independent experiments. D) Serum cytokines measured by Legendplex from mice treated as in 4C. Mixed-effects model.
Geometric Mean ± Geo SD.

